# Dysregulated neuroimmune interactions and sustained type I interferon signaling after human immunodeficiency virus type 1 infection of human iPSC derived microglia and cerebral organoids

**DOI:** 10.1101/2023.10.25.563950

**Authors:** Andrew J. Boreland, Alessandro C. Stillitano, Hsin-Ching Lin, Yara Abbo, Ronald P. Hart, Peng Jiang, Zhiping P. Pang, Arnold B. Rabson

## Abstract

Human immunodeficiency virus type-1 (HIV-1) associated neurocognitive disorder (HAND) affects up to half of HIV-1 positive patients with long term neurological consequences, including dementia. There are no effective therapeutics for HAND because the pathophysiology of HIV-1 induced glial and neuronal functional deficits in humans remains enigmatic. To bridge this knowledge gap, we established a model simulating HIV-1 infection in the central nervous system using human induced pluripotent stem cell (iPSC) derived microglia combined with sliced neocortical organoids. Upon incubation with two replication-competent macrophage-tropic HIV-1 strains (JRFL and YU2), we observed that microglia not only became productively infected but also exhibited inflammatory activation. RNA sequencing revealed a significant and sustained activation of type I interferon signaling pathways. Incorporating microglia into sliced neocortical organoids extended the effects of aberrant type I interferon signaling in a human neural context. Collectively, our results illuminate the role of persistent type I interferon signaling in HIV-1 infected microglial in a human neural model, suggesting its potential significance in the pathogenesis of HAND.

**Highlights of the work:**

- HIV-1 productively infects iPSC-derived microglia and triggers inflammatory activation.
- HIV-1 infection of microglia results in sustained type I interferon signaling.
- Microglia infected by HIV-1 incorporate into sliced neocortical organoids with persistent type I interferon signaling and disease risk gene expression.

## INTRODUCTION

The global burden of human immunodeficiency virus-1 (HIV-1) infection remains profound, with neurocognitive complications manifesting in a significant proportion of infected individuals despite the remarkable success of combined antiretroviral therapy (cART) in reducing morbidity and mortality. Up to 55% of HIV-1 positive people experience neurological complications that may develop into HIV associated neurocognitive disorder (HAND) (Simioni et al. 2010; Sacktor et al. 2016; Eggers et al. 2017; Sacktor 2018). This debilitating disorder manifests across a spectrum of cognitive, motor, affective, and behavioral impairments, which can severely impact the quality of life for affected individuals, emphasizing an urgent need for a better understanding of its underlying mechanisms (Lew et al. 2018; Israel et al. 2019). The etiology and neuropathogenesis of HAND remain enigmatic, as there has been no evidence of neuronal HIV- 1 infection. Decades of research suggest that HAND development is caused by a combination of both direct and indirect effects involving viral proteins, inflammatory cytokines and chemokines, and dysregulated neuroimmune interactions (Koenig et al. 1986; Gendelman et al. 1994; Rappaport et al. 1999; González-Scarano and Martín-García 2005; Kraft-Terry, Buch, Fox, and Gendelman 2009; Lindl, Marks, Kolson, and Jordan-Sciutto 2010; Burdo, Lackner, and Williams 2013; Alvarez-Carbonell et al. 2020). Interestingly, clinical symptoms of HAND correlate more closely with cytokine production, neuronal pathology, decreased synaptic density, and increased microglia than with central nervous system (CNS) viral load (Kaul, Garden, and Lipton 2001; Kaul et al. 2005). Currently there is a lack of mechanistic understanding due to a need for better experimental models that can address these questions, especially regarding neuroimmune interactions.

A pivotal component of neuroimmune interactions is the innate immune response, including the production and signaling of interferons, which have recently been implicated in the neuropathology of various neurological diseases [Reviewed in (Hofer and Campbell 2013; McGlasson, Jury, Jackson, and Hunt 2015)]. Interferons and their downstream signaling through interferon stimulated genes (ISGs) are primarily an innate immune response to viral infection [reviewed in (Platanias 2005)]. However, recent literature suggests a role for dysregulated type I interferon signaling in neurological diseases such as Alzheimer’s disease (Roy et al. 2020; Roy et al. 2022), COVID-19 related brain fog (Lee and Shin 2020; Phetsouphanh et al. 2022), aging associated dementia (Baruch et al. 2014), and HIV associated dementia (Rho et al. 1995; Krivine et al. 1999; Perrella et al. 2001; Anderson et al. 2017; Solomon et al. 2020). Clinically, type I interferons have been shown to be a driver of cognitive impairment both in diseased states and when administered exogenously (Merimsky, Reider-Groswasser, Inbar, and Chaitchik 1990). Indeed, many studies using interferon treatment for various severe cancers and chronic viral hepatitis C have reported that patients receiving interferon-alpha displayed cognitive, mood, and behavioral deficits that arose and worsened with prolonged, multiweek treatment regimens (Christina, Randall, and Arthur 1991; Pavol et al. 1995; Capuron, Ravaud, and Dantzer 2001; Capuron et al. 2002; Wichers et al. 2005; Raison, Demetrashvili, Capuron, and Miller 2005). Taken together, these studies indicate a role for aberrant type I interferon signaling in compromising neuronal and glial function, leading to cognitive deficits and degeneration.

Microglia, as the resident immune cells of the CNS and the primary CNS target and reservoir of HIV-1 infection (Tang et al. 2023), contribute significantly to the pathophysiology of HAND, largely through their secretory and functional dysregulation. Under homeostatic conditions, microglia play important roles in synaptic pruning, phagocytic functions, immune surveillance, and extracellular matrix remodeling (Paolicelli et al. 2011; Casano and Peri 2015). When challenged, microglial activation can be either neurotoxic, producing proinflammatory cytokines and chemokines, or neurotrophic, producing anti-inflammatory cytokines (Durafourt et al. 2012). Microglia exhibit a heterogeneous spectrum of activation and disease-associated states with unique properties and consequences on neuronal functions (Gosselin et al. 2017; Song and Colonna 2018; Hammond et al. 2019).

Since HIV-1 specifically infects humans, the ability of traditional animal model organisms to replicate the complex disease processes of HAND is inherently limited. HIV-1-Simian immunodeficiency virus chimeras (SHIVs) have been studied in monkey models. However, these models are limited in their availability and utility for molecular analyses [reviewed in (Byrnes et al. 2022; Moretti et al. 2021)]. Recent advances in stem cell technologies have opened up new possibilities for modeling and studying HAND, particularly the development of human iPSC derived neurons, astrocytes, and microglia capable of mimicking complex in vivo neuroimmune interactions when grown in three dimensional cerebral organoid cultures (Abud et al. 2017; Ormel et al. 2018; Xu et al. 2021). Cerebral organoids offer a powerful and physiologically relevant platform to study complex neuroimmune interactions, but traditional organoid differentiation protocols do not produce microglia [reviewed in(Dos Reis, Sant, and Ayyavoo 2023; Swingler et al. 2023; Wei et al. 2023)]. Recent studies have demonstrated the feasibility of infecting iPSC-derived macrophages (Vaughan-Jackson et al. 2021) and microglia (Rai et al. 2020) with HIV-1. Moreover, these infected microglia have been effectively cocultured in various model systems, including human stem cell derived neural tricultures (Alvarez-Carbonell et al. 2020; Ryan et al. 2020), primary neuron progenitor cell derived organoids (dos Reis et al. 2020), iPSC-derived cerebral organoids (Gumbs et al. 2022), and a humanized mouse model (Min et al. 2023).

In this study, we hypothesized that by infecting iPSC-derived microglia with replication competent HIV-1 and integrating them into sliced neocortical organoids, we could recapitulate neuroimmune dysregulation related to HAND pathology. We demonstrate that HIV-1 infection of iPSC-derived microglia leads to persistent activation of type I interferon signaling and microglial activation in both microglial monocultures and in the context of cerebral organoids.

## RESULTS

### Human microglia are efficiently generated from iPSCs

To induce microglial differentiation, we used three iPSC lines and employed previous protocols (Haenseler et al. 2017; Xu et al. 2021). To mimic in vivo microglial development from the yolk sac, iPSC embryoid bodies were patterned toward a mesodermal yolk sac lineage that generates primitive macrophage precursors (PMPs) after 3 weeks of culture (**Figure 1A**). The identity of the PMPs was confirmed by immunocytochemistry against CD235, a marker for yolk sac (Claes, Van den Daele, and Verfaillie 2018), and CD43, a marker for hematopoietic progenitor like cells (Vodyanik, Thomson, and Slukvin 2006) (**Figure 1B**). Upon further maturation under the influence of a colony stimulating factor 1 receptor (CSF1R) antagonist, interleukin 34 (IL-34), and granulocyte maturation factor (GM-CSF), PMPs exhibit a more pronounced microglial phenotype. This was evidenced by a significant increase in mRNA expression of the microglial markers ionized calcium binding molecule 1 (IBA1) and purinergic receptor P2Y12 across all three cell lines (**Figure 1C**, biological replicates of each cell line shown in **Figure S1**). Supporting these gene expression results, we noted robust immunocytochemical staining of the microglial markers IBA1, P2Y12, CX3CR1, transmembrane protein 119 (TMEM119), and transcription factor PU.1 in maturing microglia from these same lines (**Figure 1D**). Upon coculture of microglia with iPSC-derived neurons (Xu et al. 2021), we found that microglia display a more elaborate morphology (**Figure 1E**). Furthermore, costaining with the neuronal dendritic marker microtubule-associated protein 2 (MAP2) and the presynaptic marker synapsin 1 (SYN1) revealed that microglial processes interact with dendrites and colocalize with synaptic puncta, indicating putative microglia-synapse interactions (**Figure 1F**).

**Figure 1:**
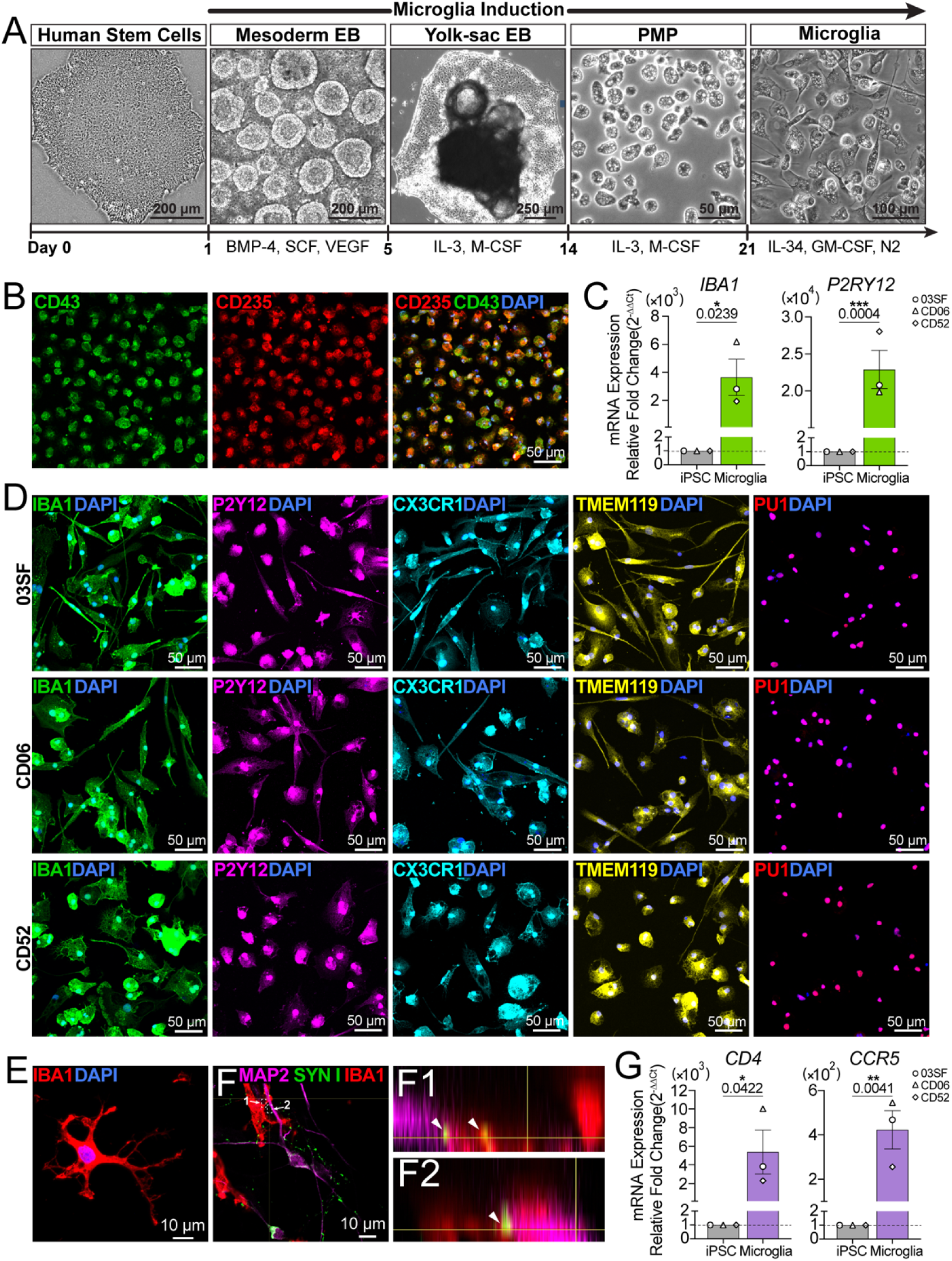
Human microglia are efficiently generated from induced pluripotent stem cells. (A) Representative brightfield images showing the induction of iPSCs to mesodermal yolk sac embryoid bodies (EB) and then to primitive microglia progenitor cells (PMPs) and microglia. (B) Representative confocal images of PMPs expressing the yolk sac marker CD235 and the hematopoietic progenitor cell marker CD43 in the 03SF cell line. (C) RT-qPCR analysis of *IBA1* and *P2RY12* mRNA expression in iPSCs and iPSC-derived microglia; n=3 cell lines (03SF, CD06, CD52); t test; data are presented as the mean ± SEM. (D) Representative confocal images of microglia stained for the microglial markers IBA1, P2Y12, CX3CR1, TMEM119, and PU1 and costained with DAPI; n = 3 cell lines (03SF, CD06, CD52). (E) Representative confocal image of a microglial cell displaying ramified morphology after 4 weeks of coculture with neurons. (F) Representative confocal images of a 03SF cell line iPSC-derived microglial neuronal coculture stained for neurons (MAP2), synapses (SYN1), and microglia (IBA1). (F1 and F2) Orthogonal view of microglial (IBA1+) processes colocalized with neuronal dendrites (MAP2) and synapses (SYN1). (G) RT-qPCR analysis of *CD4* and *CCR5* mRNA from iPSCs and microglia; n=3 cell lines (03SF, CD06, CD52); t test; data are presented as the mean ± SEM.

Given the tropism of HIV-1 for CD4 receptor expressing cells and the requirement of CCR5 coreceptor expression for infection of monocyte/macrophage related cell lineages, we assessed the gene expression of *CD4* and *CCR5* in iPSC-derived microglia. Results showed significantly increased mRNA expression of *CD4* and *CCR5* in microglia differentiated from all three iPSC lines (**Figure 1G**, biological replicates of each cell line shown in **Figure S1**). These results demonstrate that microglia can be successfully and reproducibly generated from iPSCs. Crucially, these cells express appropriate markers for microglial identity and infection by HIV-1.

### iPSC derived microglia are productively infected by HIV-1 and exhibit inflammatory activation

Having confirmed the generation and maturation of iPSC-derived microglia, we next determined whether these cells could sustain a productive infection by replication competent, macrophage tropic HIV-1. To this end, we incubated male 03SF iPSC-derived microglia with JRFL HIV (2 ng/ml), an R5 tropic HIV-1 strain isolated from the cerebral spinal fluid (CSF) of a patient with severe HAND (Koyanagi et al. 1987; Merrill et al. 1992). Compared to uninfected controls, HIV- 1-infected microglia robustly expressed HIV-1 *gag/pol* RNA within 6-9 days after infection (Yu et al. 1996) (**Figure 2A**). As a key indicator of productive HIV-1 infection, the production of secreted p24 was measured (Chargelegue and O’Toole 1992; Marodon, Warren, Filomio, and Posnett 1999). Using ELISA to detect secreted p24 in the culture supernatant, we observed increased protein production starting 1 week post infection, peaking around day 16, and then maintaining consistent levels for more than 2 weeks (**Figure 2B**). Finally, we found that after just 4 days post infection, circular polymerase chain reaction (PCR) products from both long terminal repeats (LTR) were detected in HIV-1 incubated cultures indicating the presence of active viral replication (Butler, Hansen, and Bushman 2001) (**Figure S2**).

**Figure 2:**
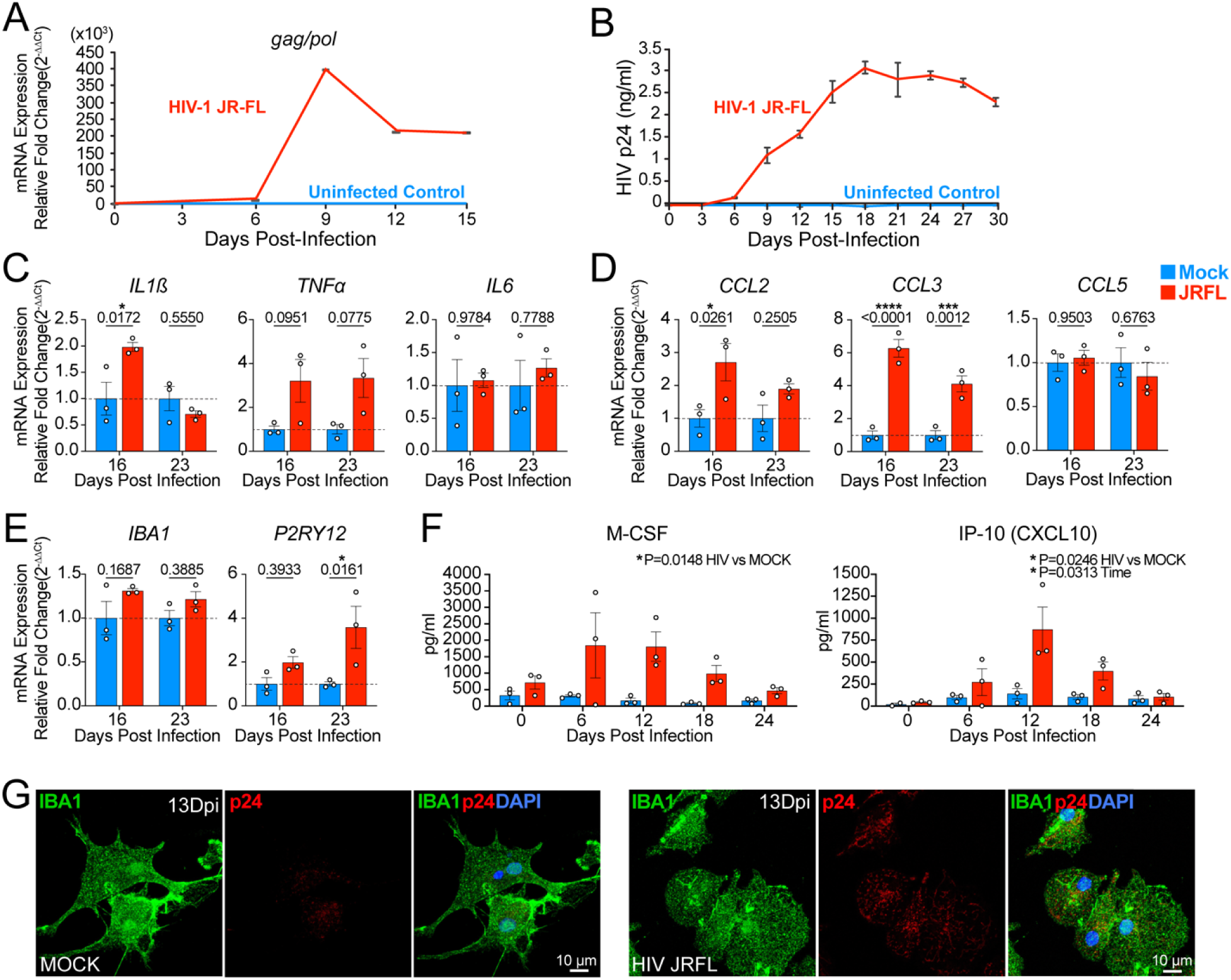
HIV-1-infected microglia exhibit productive infection, activation, and dysregulated cytokine/chemokine production. (A) RT-qPCR analysis of viral *gag/pol* mRNA after infection with HIV-1 JRFL in microglia monocultures; n=3 independent infections of 03SF line iPSC-derived microglia. (B) ELISA analysis of secreted p24 in the culture supernatant of microglia infected with HIV-1 JR-FL reveals productive infection; n=3 independent infections of 03SF iPSC-derived microglia. (C) RT-qPCR analysis of *IL1*β*, TNF*α, and *IL-6* mRNA at days 16 and 23 post infection with HIV-1 JRFL; n=3 independent infections of 03SF iPSC-derived microglia, t test; data are presented as the mean ± SEM. (D) RT-qPCR analysis of *CCL2, CCL3*, and *CCL5* mRNA at days 16 and 23 post infection with HIV-1 JRFL; n=3 independent infections of 03SF iPSC-derived microglia. (E) RT-qPCR analysis of *IBA1* and *P2RY12* mRNA at days 16 and 23 post infection with HIV-1 JRFL; n=3 independent infections of 03SF iPSC-derived microglia. (F) Luminex cytokine/chemokine analysis of secreted M-CSF and IP-10/CXCL10 culture supernatant from HIV-1 JRFL-infected microglia; n=3 independent infections of 03SF microglia, two-way ANOVA; data are presented as the mean ± SEM. (G) Representative confocal images of nucleocapsid protein p24 staining in HIV-1 JRFL-infected 03SF- derived microglia (IBA1^+^).

After establishing the infectability of iPSC microglia with HIV-1 JRFL, we then asked whether HIV-1 infection elicits differential expression of inflammatory and canonical microglial mRNAs using reverse transcription[quantitative PCR (RT-qPCR). In this second experiment found we found HIV-1 infection augmented the expression of the inflammation-associated genes interleukin 1ß (IL1ß, on day 16) and tumor necrosis factor alpha (TNFα), while the expression of interleukin 6 (IL6) displayed no change (**Figure 2C**). We then examined the mRNA expression of the chemokines CCL2 (monocyte chemoattractant protein 1, MCP-1), CCL3 (macrophage inflammatory protein 1 alpha, MIP-1), and CCL5 (RANTES). These three chemokines play important roles in immune cell recruitment and migration to sites of inflammation and infection and have been found to be elevated in cognitively impaired individuals with HIV-1 (Letendre, Lanier, and McCutchan 1999). A significant elevation in the mRNA expression of CCL2 and CCL3 was observed by day 16 post-infection, whereas the expression of CCL5 remained unchanged (**Figure 2D**). To assess whether microglial identity was altered by HIV-1 infection, we assessed the mRNA expression of the canonical microglial genes IBA1 and P2Y12. While IBA1 remained stable, P2Y12 exhibited significantly increased expression by day 23 post infection (**Figure 2E**). Next, we asked how HIV-1 infection might alter microglial chemokine and cytokine production. Although the levels of most cytokines were below the detection threshold (**Table S1**), we noted a significant surge in the secretion of chemokines M-CSF and interferon inducible protein 10 (IP-10/CXCL10) following HIV-1 infection in the same supernatant used in figure 2B (**Figure 2F**). Finally, to visualize HIV-1 infected cells, we detected HIV-1 p24 protein using a biotin-streptavidin amplification protocol (Eugenin and Berman 2016). On day 13 post infection with JRFL, p24 positive microglia were found (**Figure 2G**). Collectively, our findings suggest that iPSC-derived microglia can sustain an R5 tropic HIV-1 infection and exhibit an inflammatory activation profile consistent with previous studies.

To confirm that HIV-1 JRFL infection of human iPSC-derived microglia resulted in productive infection and inflammatory activation, we chose a female iPSC line, CD06, and infected it with the JRFL strain as well as a second R5 tropic HIV-1 strain, YU2 (Li et al. 1991), isolated from the CSF of a different patient with severe HAND. We confirmed infection by RT-qPCR of viral *gag-pol* RNA, which exhibited significantly increased expression for both strains by day 16 post-infection, with a subsequent decrease (**Figure 3A**). Using ELISA to detect secreted p24 revealed productive infection for both viral strains (**Figure 3B**). We then assessed the expression of key inflammatory markers and microglial core mRNAs by qPCR. Both JRFL and YU2 infections led to a significant increase in IL1β expression compared to uninfected microglia (**Figure 3C**). Similarly, TNFα was significantly upregulated in the first two weeks of infection but decreased as the productive infection waned (**Figure 3C**). We also observed increased expression of the chemokines CCL2 and CCL3 by days 16 and 23 post infection with both strains (**Figure 3D**). This upregulation then decreased by day 27, correlating with the reduction of productive infection as indicated by the p24 ELISA (**Figure 3A**). In comparison to the unaltered IBA1 expression we found previously, HIV-1 led to a significant reduction in IBA1 expression in CD06 infected microglia (**Figure 3E**). Similar to 03SF infected microglia, CD06 had significant increased expression of P2RY12 (**Figure 3E**). Together, these results demonstrate infectability of human iPSC-derived microglia with multiple, macrophage tropic HIV-1 strains and their ability to elicit an inflammatory response.

**Figure 3:**
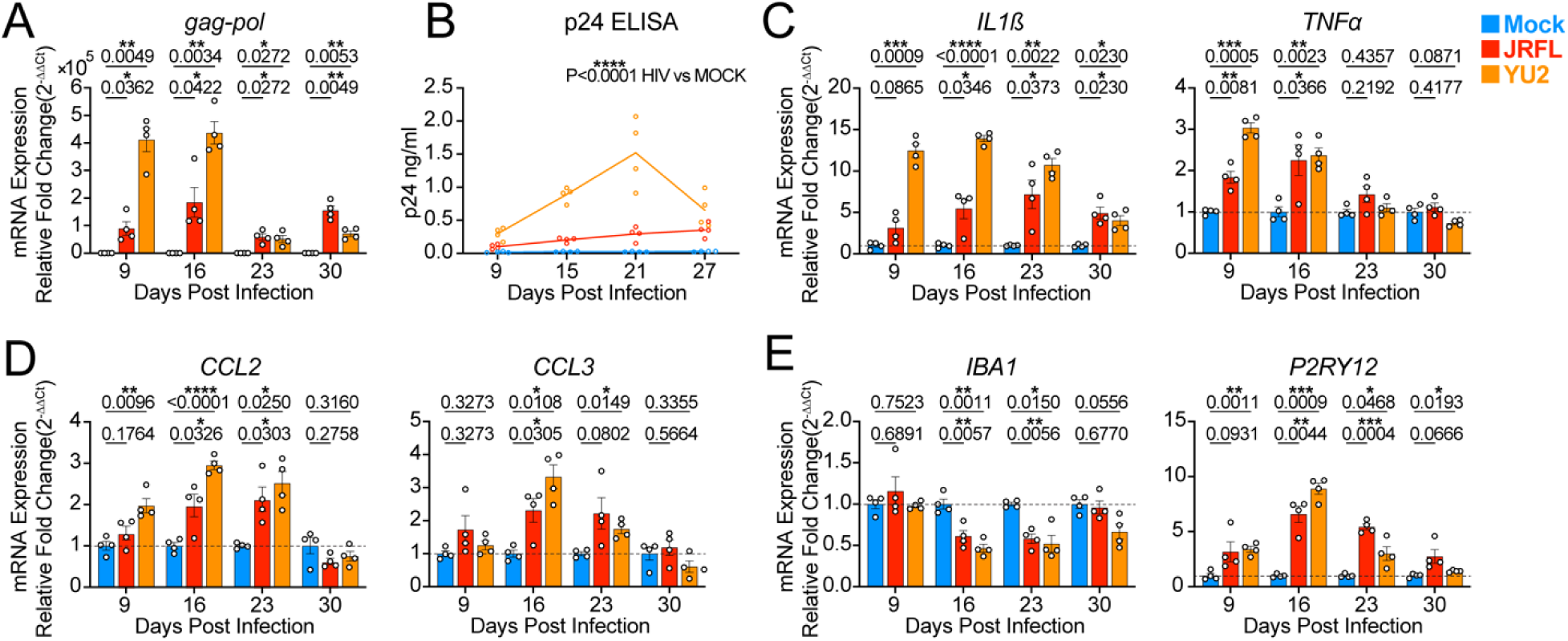
HIV-1-infected microglia exhibit productive infection, activation, and dysregulated cytokine/chemokine production. (A) RT-qPCR analysis of viral *gag/pol* mRNA at days 9, 16, 23, and 30 post infection with HIV-1 JRFL and HIV-1 YU2; n=3 independent infections of CD06 iPSC-derived microglia. t test (B) ELISA analysis of secreted p24 in the culture supernatant of microglia infected with HIV-1 JRFL and HIV-1 YU2 at days 9, 15, 21, and 27 reveals productive infection; n=3 independent infections of CD06 iPSC-derived microglia. (C) RT-qPCR analysis of *IL1*β and *TNF*α mRNA at days 9, 16, 23, and 30 post infection with HIV-1 JRFL and HIV-1 YU2; n=3 independent infections of CD06 iPSC-derived microglia. (D) RT-qPCR analysis of *CCL2* and *CCL3* mRNA at days 9, 16, 23, and 30 post infection with HIV-1 JRFL and HIV-1 YU2; n=3 independent infections of CD06 iPSC-derived microglia. (E) RT-qPCR analysis of *IBA1* and *P2RY12* mRNA at days 9, 16, 23, and 30 post infection with HIV-1 JRFL and HIV-1 YU2; n=3 independent infections of CD06 iPSC-derived microglia.

### Type I interferon stimulated gene expression and sustained activation following HIV-1 infection

To investigate the transcriptional landscape of HIV-1 infected microglia, we performed bulk RNA sequencing on male 03SF iPSC-derived microglia infected by HIV-1 JRFL day 16 post infection. Principal component analysis (PCA) revealed clear separation of the infected vs mock infected samples (**Figure S2**). Differential gene analysis revealed differentially expressed genes (DEGs) including several classic ISGs, such as IFIT3, MX2, ISG15, RSAD2 and others (**Figure 4A**) (**Table S2**). Gene set enrichment analysis (GSEA) confirmed a significant enrichment of genes related to the interferon alpha response (**Table S3**) (**Figure 4B**). Gene ontology analysis identified activation of innate immune responses and several related terms in the biological process component (**Figure 4C, D** and **Figure S3**). The upregulated DEG GO set includes pathways falling into the overall signaling clusters related to interferon signaling and positive regulation of antiviral biotic, interferon alpha/beta, defense against virus, cytokine mediated and interferon mediated pathways, and negative regulation of viral processes. Among the downregulated DEG GO clusters are pathways related to positive regulation of antigen processing, extracellular organization, granulocyte/leukocyte migration, chemokine-mediated signaling, and spindle assembly checkpoints. Importantly, HIV-1 infection was found to lead to significant induction of many ISGs, including ISG15, IRF7, MX1, OAS1, OAS2, OAS3, STAT1, CXCL10, and IFITM3 (**Figure 4A, E**). **Figure 4F** shows the expression of various genes encoding SIGLEC proteins that are responsible for regulating microglial cell[cell interactions [Reviewed in (Crocker, Paulson, and Varki 2007)]. Consistent with previous studies, we found that HIV-1 infection causes significant upregulation of SIGLEC1, which has been implicated in the direct cell[cell transmission of HIV (Izquierdo-Useros et al. 2012; Zou et al. 2011). However, we found that SIGLEC11 and SIGLEC16 were significantly downregulated after infection.

**Figure 4:**
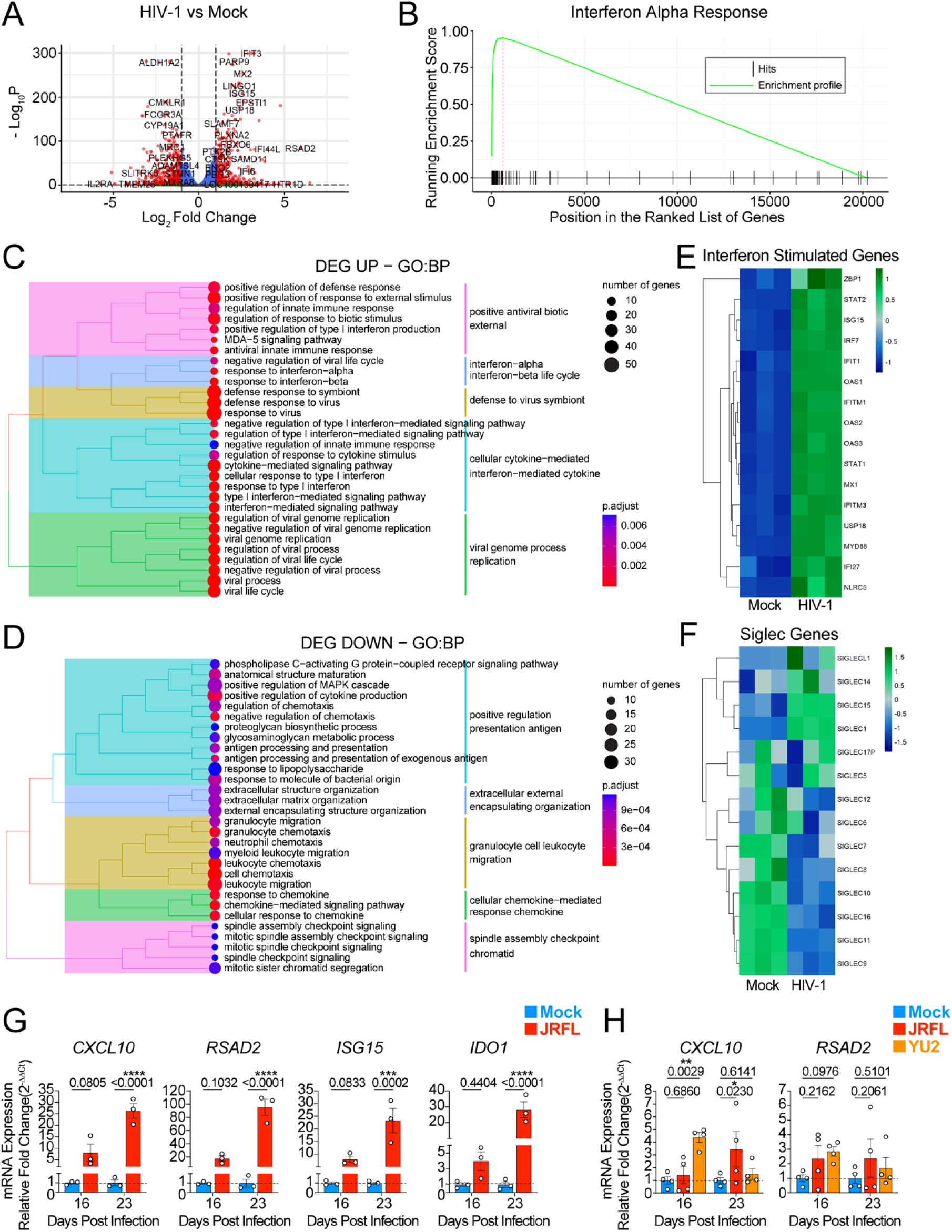
HIV-1 causes enrichment of type I interferon signaling pathways and sustained interferon activation. (A) A volcano plot illustrating differentially expressed genes (DEGs) in HIV-1 JRFL-infected vs mock- infected microglia on day 16 post infection; n=3 independent infections of 03SF line iPSC-derived microglia. (B) GSEA plot showing enrichment of IFNα-responsive genes in HIV-1 JRFL-infected vs mock-infected microglia on day 16 post infection (NES, normalized enrichment score; FDR, false discovery rate). (C) Gene Ontology (GO) analysis of upregulated DEGs in HIV-1 JRFL vs mock-infected microglia on day 16 post infection; n=3 independent infections of 03SF microglia. (D) Gene Ontology (GO) analysis of downregulated DEGs in HIV-1 JRFL vs mock-infected microglia on day 16 post infection; n=3 independent infections of 03SF microglia. (E) A heatmap illustrating significant upregulation of interferon-stimulated genes in HIV-1-infected vs mock-infected microglia on day 16 post infection; n=3 independent infections of 03SF microglia. (F) A heatmap illustrating SIGLEC gene expression in HIV-1-infected vs mock-infected microglia on day 16 post infection; n=3 independent infections of 03SF microglia. (G) RT-qPCR analysis of *CXCL10*, *RSAD2*, *ISG15*, and *IDO1* mRNA from HIV-1 JRFL-infected vs mock-infected microglia at days 16 and 23 post infection; n = 3 independent infections of 03SF iPSC- derived microglia, t test; *p < 0.05, **p < 0.01, ***p < 0.001; data are presented as the mean ± SEM. (H) RT-qPCR analysis of *CXCL10* and *RSAD2* mRNA from HIV-1 JRFL- and HIV-1 YU2-infected and mock-infected microglia at days 16 and 23 post infection; n = 4 independent infections of CD06 iPSC- derived microglia, t test; *p < 0.05, **p < 0.01, ***p < 0.001; data are presented as the mean ± SEM.

We then validated the expression of a subset of the ISGs identified in the sequencing dataset by RT-qPCR of JRFL infected 03SF iPSC-derived microglia. We found significant upregulation of CXCL10, RSAD2, ISG15, and IDO1 (**Figure 4G**). Elevated expression of these genes persisted over an additional week of infection on day 23 (**Figure 4G**). Additionally, we performed RT-qPCR on CD06 iPSC-derived microglia infected with either JRFL or YU2 HIV strains. We also found increased expression of CXCL10 and RSAD2 in these HIV-infected cells; however, only the day 16 YU2 condition reached significance (**Figure 4H**). Comparison of Figure 4G and 4H indicates potential differences with infecting different cell lines. Together, these data support the conclusion that HIV-1 infection induces sustained type I interferon signaling in iPSC-derived microglia and alters the gene regulation of many microglial biological processes including interferon signaling.

### HIV-1 infected microglia incorporate into cerebral organoids

After characterizing the response of isolated human iPSC-derived microglia to HIV-1 infection we then investigated whether HIV-1 infected microglia could incorporate into iPSC-derived cerebral organoids and whether interferon signaling was similarly upregulated in microglia and potentially other neural cell types. Cerebral organoids were grown using modified, published protocols (Lancaster and Knoblich 2014; Paşca et al. 2015; Sloan et al. 2018; Yoon et al. 2019; Qian et al. 2020; Xu et al. 2021). To create microglia-containing organoids, we cocultured human microglia with sliced neocortical organoids (SNO), prepared as described previously (Qian et al. 2020). SNOs exhibit enhanced neurogenesis and laminar expansion (Qian et al. 2020). We found that microglia exhibit the ability to incorporate with organoid slices to form human microglia containing sliced neocortical organoids (mSNOs). **Figure 5A** shows mSNO containing neurons (MAP2^+^), astrocytes (GFAP^+^), and microglia (IBA1^+^). These neuroimmune organoids provide a model system for studying the impact of HIV-1 infection on the physiology, localization, and interactions of these three cell types.

**Figure 5:**
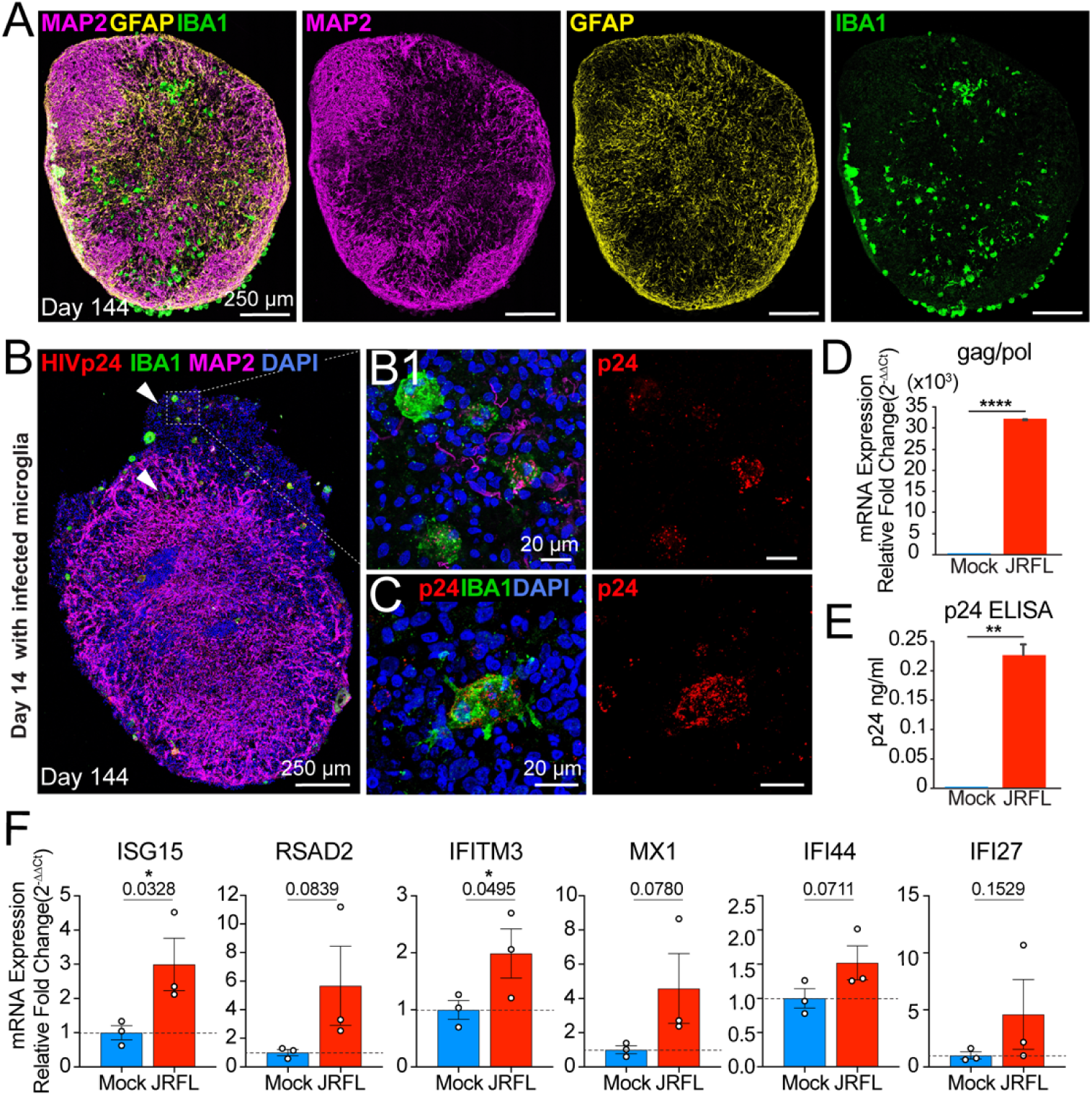
HIV-1-infected microglia incorporate into sliced cerebral organoids. (**A**) Representative confocal images of a day 144 organoid with neurons (MAP2^+^), astrocytes (GFAP^+^), and microglia (IBA1^+^). (B) Representative confocal images of day 144 organoids 14 days after the addition of HIV-1 JRFL- infected or mock-infected microglia; (B1) shows a high-magnification image of HIV-1-infected microglia from (B). (C) High-magnification image of a single HIV-1-infected microglial cell within a sliced organoid. (D) RT-qPCR of *gag/pol* mRNA from HIV-1 JRFL-infected or mock-infected microglia organoids on day 14; n=3 independent infections of 03SF microglia; one-tailed t test; ****P<0.001; error bars represent SEM. (E) ELISA analysis of secreted p24 in culture supernatant from HIV-1 JRFL-infected or mock-infected microglia organoid day 14; n=3 independent infections of 03SF microglia; one-tailed t test; **p<0.01; data are presented as the mean ± SEM. (F) RT-qPCR analysis of *ISG15*, *RSAD2*, *IFITM3*, *MX1*, *IFI44*, and *IFI27* mRNA from HIV-1 JRFL-infected or mock-infected microglia organoids on day 14; n=3 independent infections of 03SF microglia; one-tailed t test; *p < 0.05, **p < 0.01, ***p < 0.001; data are presented as the mean ± SEM.

To create an HIV-1 infected human cerebral organoid, we infected 03SF iPSC-derived microglia with JRFL HIV. Following 2 weeks in culture, infected or noninfected microglia were added to 400 µM organoid slices and cultured for 2 additional weeks. On day 14 post coculture with microglia, mSNOs were fixed for immunohistochemistry or total RNA was extracted for RT-qPCR analysis. HIV-1-infected microglia (p24^+^, IBA1^+^) were detected in a mSNO (**Figure 5B&C**). Interestingly, neurons (MAP2^+^) appeared to be sparser in regions of HIV-1-infected microglia (**Figure 5B** “arrows”). These regions had scattered MAP2^+^ processes (**Figure 5B1**) compared with the denser MAP2 staining throughout the bulk of the organoid. HIV-1 *gag/pol* RNA exhibited significant expression in mSNOs containing HIV-1 infected microglia (**Figure 5D**). To provide evidence of productive infection in mSNOs, we identified secreted p24 only in the media of organoid cultures that had received HIV-1 infected microglia (**Figure 5E**).

Because we observed significant and sustained interferon signaling in microglial monocultures, we evaluated mRNA expression of a panel of ISGs (ISG15, RSAD2, IFITM3, MX1, IFI44, IFI27). We found that ISG15 mRNA was significantly upregulated and RSAD2 also showed an increasing trend of expression in mSNOs containing HIV-1 infected microglia (**Figure 5F**). IFITM3 has been touted as a marker for interferon-responsive microglia and has been shown to play a newly discovered role in amyloid precursor protein processing to amyloid beta (Hur et al. 2020). Interestingly, we found significant upregulation of IFITM3 mRNA in mSNOs containing HIV-1 infected microglia (**Figure 5F).** We also evaluated the expression of MX1, IFI27, and IFI44, which, along with ISG15 and RSAD2, were among the top 10 persistently upregulated genes in a recent study on two cohorts of 22 HIV-1 seroconverts from Uganda and 22 from South Africa (Mackelprang et al. 2023). Expression of MX1, IFI27, and IFI44 appeared to be increased but failed to reach significance. Especially considering the small fraction of HIV-1 infected microglia in the cultured mSNO (**Figure 5B**), these increases suggest that HIV-1 infected microglia are associated with increased interferon signaling in the organoids and may have indirect consequences for neuronal viability and inflammatory responses in other cell types.

## DISCUSSION

In this study, we used iPSC-derived human microglia and microglia containing cortical organoids to model HIV-1 related neuroimmune dysregulation and pathology. In this model, iPSC-derived microglia were shown to be infected with two strains of replication competent HIV- 1, JRFL and YU2, and exhibited inflammatory activation, as shown by gene expression changes and cytokine/chemokine secretion (**Figures 2 & 3**). Importantly, we found persistent activation of type I interferon response pathways and downstream interferon stimulated genes (ISGs), which may have important implications for understanding the pathogenesis of HAND.

In comparison to the recently developed primary human neural progenitor cell-derived organoid HIV-1 model (dos Reis et al. 2020) and the 2D neuron, astrocyte, and microglia “triculture” HIV-1 model (Ryan et al. 2020), an iPSC-derived human microglia containing sliced neocortical organoid model has distinct advantages. Organoids recapitulate 3D cell-cell and cell-matrix interactions more faithfully than 2D cultures and develop complex tissue architecture (Lancaster et al. 2013; Paşca et al. 2015; Birey et al. 2017; Quadrato et al. 2017; Velasco et al. 2019); thus, organoids are well suited to studying aspects of altered brain anatomy and physiology seen in individuals with *HAND* (Di Sclafani et al. 1997; Clifford et al. 2017; Israel et al. 2019). While others have also generated microglia containing organoids using a protocol in which microglia innately develop within organoids, this system is associated with a high degree of heterogeneous cell types (Gumbs et al. 2022), which could make dissection of the relative contributions and effects on specific cell types more complex. In our method, we have greater freedom to manipulate the timing, proportion, and infection stage of microglia addition in sliced neocortical organoids. While some previous studies used pseudotyped virus (dos Reis et al. 2020), our study examined infection using two macrophage tropic, replication competent strains of HIV-1. Our findings, showing induction of inflammatory gene expression associated with active viral expression align with recent studies on brain samples from HIV-1^+^ patients from the National NeuroAIDS Tissue Consortium (NNTC). In that study, HIV-1 RNA load was found to correlate with increased interferon signaling and inflammatory gene sets while inversely correlating with neuronal and synaptic gene sets, suggesting neuronal pathology (Sanna et al. 2021).

Microglia secrete diverse cytokines and chemokines implicated in HIV-1 neuropathology (Hanisch 2002; Ramesh, MacLean, and Philipp 2013; Chen et al. 2017). Cytokine/chemokine analysis of the supernatant of HIV-1 infected microglia identified the secretion of M-CSF and IP- 10/CXCL10 (**Figure 2F**). M-CSF overexpression has been linked to microglial proliferation, increased phagocytosis, cytokine secretion, and inflammatory responses in both Alzheimer’s disease (AD) and HAND, as well as increased microglial susceptibility to HIV-1 infection and increased virus replication (Gallo et al. 1990; Kalter et al. 1991; Kutza et al. 2000; Mitrasinovic et al. 2001; Kutza, Fields, Grimm, and Clouse 2002; Smith et al. 2013). IP-10/CXCL10 is a chemotactic chemokine that recruits leukocytes, T cells, and monocytes while also being neurotoxic and strongly correlated with HIV-1 related disease progression (van Marle et al. 2004; Jiao et al. 2012; Lei, Yin, Shang, and Jiang 2019). Clinical studies have shown elevated IP-10/CXCL10 levels (∼200-1000 pg/ml) in the cerebral spinal fluid of HIV-1^+^ individuals (Kolb et al. 1999; Peluso et al. 2017); these levels are strikingly similar to our observations (**Figure 2F**). CXCL10/IP-10 has been shown to be induced by HIV-1 proteins gp120, Nef, and Tat (Kutsch, Oh, Nath, and Benveniste 2000; Asensio et al. 2001; van Marle et al. 2004; Lei, Yin, Shang, and Jiang 2019). Importantly, IP-10/CXCL10 and RANTES (CCL5) were the strongest predictors of neurocognitive impairment in a screen of 139 HIV-1^+^ individuals (Ruhanya et al. 2021), similar to findings from earlier studies (Valverde-Villegas et al. 2018; Burlacu et al. 2020). HIV-1 brain infection leads to an abnormal inflammatory state and dysregulated cytokine/chemokine production (Kaul, Garden, and Lipton 2001; González-Scarano and Martín-García 2005; Burdo, Lackner, and Williams 2013). Despite this knowledge, there remains a large gap in our understanding of how CXCL10 and other cytokines/chemokines associated with CNS HIV-1 infection alter aspects of neuronal excitability, synaptic transmission, and inward and outward currents vital to healthy neuronal physiology.

A striking finding in our study was the persistent activation of interferon response genes. Induction of these pathways is known to be a hallmark of HIV-1 infection in monocytes and macrophages, much more prominently than in T cells (Berg et al. 2014; Aso, Ito, Koyanagi, and Sato 2019; Elsner et al. 2020; Jeremiah et al. 2023). Thus, it is not surprising to have seen this strong response in microglia, which share many physiologic properties with macrophages [reviewed in (Li and Barres 2018)]. Most studies suggest that the interferon response is relatively transient; thus, the persistent expression of at least a subset of ISGs may be a relatively unique feature of microglial HIV-1 infection that may have important pathophysiologic consequences. Increasing evidence suggests a role for type I interferons in cognitive-related deficits seen in patients with various neurodegenerative diseases. Using single-cell sequencing, researchers have dissected a unique sub-population of interferon-responsive microglia that may potentiate dysregulated immune interactions in the form of aberrant pruning of synapses, altered phagocytic function, and dysregulated cytokine/chemokine production (Jin et al. 2022). Constitutive overexpression of type I interferons seen in the “interferonopathies” can be associated with profound CNS disease [reviewed in (Goldmann, Blank, and Prinz 2016), and (Blank and Prinz 2017)](Wiley, Steinman, and Wang 2023). Patients receiving type I interferons therapeutically may experience cognitive impairment and neurological symptoms (Christina, Randall, and Arthur 1991; Pavol et al. 1995; Capuron, Ravaud, and Dantzer 2001; Capuron et al. 2002; Wichers et al. 2005; Raison, Demetrashvili, Capuron, and Miller 2005).

Whether interferon induction inhibits or enhances HIV infection and latency in microglia remains to be understood. Recent studies have highlighted that interferon can inhibit HIV infection of primary microglia but less so of iPSC-derived microglia, as well as macrophages (Nasr et al. 2017; Woodburn et al. 2022). Interestingly, interferon may promote latency in macrophages (Dickey, Martins, Planelles, and Hanley 2022). An important feature of HIV infection of microglia identified in previous in vitro studies has been the rapid induction of viral latency (Alvarez-Carbonell et al. 2019; Sreeram et al. 2022). Our cerebral organoid model will provide an interesting system to examine the induction of HIV latency, its reactivation, and its effects on both adjacent and more distant neural cells.

An important future direction, facilitated by the development of this model system, will be experiments aimed at understanding the functional consequences that HIV infection, dysregulated interferon signaling, inflammatory activity, and changes in other important genes expressed in the CNS, may have on neuronal synaptic transmission. A number of ISGs have predicted downstream effects that could trigger or enhance neurodegeneration. For example, indoleamine 2,3-dioxygenase 1 (IDO) is a key enzyme in tryptophan catabolism, leading to the formation of kynurenic acid, an antagonist of NMDA receptors (Schwarcz and Stone 2017). This pathway and its products, such as the neurotoxin and NMDAR antagonist quinolinic acid, have been previously implicated in neurodegenerative diseases, including Huntington’s, Alzheimer’s and Parkinson’s disease (Malpass 2011; Maddison and Giorgini 2015).

Overall, the HIV-1 mSNO model presents an effective in vitro platform for investigating the underlying molecular and cellular mechanisms associated with HIV-1 and human neural cell interactions. **Our results predict that interferon stimulated genes likely play a role in altering neuronal function and contributing to HAND.**

## Supporting information

Supplemental Table 1 - Luminex Cytokine Profiling

Supplemental Table 2 - deseq2 HIV vs Control anno

Supplemental Table 3 - HIV GSEA result table

Supplemental Table 4 - HIV GO and KEGG tables

## ACKNOWLEDGEMENTS

This work was supported in part by grants from the NIH (R21NS120806), a Rutgers Busch Biomedical Grant, a Rutgers Brain Health Institute Pilot Grant, and the Robert Wood Johnson Foundation (Grant #74260). AJB was supported by NIH NIGMS T32GM8339 and NIH NCATS TL1TR003019.

## AUTHOR CONTRIBUTIONS

AJB, ZPP, PJ, RPH, and ABR conceived the study. AJB designed the experiments and interpreted the data. AJB performed most of the experiments with technical assistance from ACS, HL, and YA. ACS and AJB performed qPCR. HL and AJB performed ELISA. AJB performed immunocytochemistry, immunohistochemistry, and confocal imaging. AJB and HS performed viral infections. RPH and AJB performed computational analysis of RNA-seq data. AJB prepared the figures and wrote the manuscript with input from all coauthors.

## METHODS

### Human induced pluripotent stem cell maintenance

Three human iPSC lines were used in this study (male 03SF iPSC line; female CD06 iPSC line; male CD52 iPSC line). Human iPSCs were cultured in stem cell medium consisting of MACs iPS-Brew XF complete media. The culture medium was changed daily. Human iPSCs were passaged one to two times per week onto new plates coated with Matrigel. Human iPSCs were detached with Accutase treatment for 5 min and centrifuged at 120 rcf for 5 min. The pellet was resuspended in stem cell medium plus Y-27632 (10 µM). The iPSCs used in this study were below passage 50. All studies were performed with protocols approved by Rutgers University.

### Generation of microglia

Primitive microglia precursors (PMPs) were generated from 3 iPSC cell lines using a previously established protocol (Haenseler et al. 2017). iPSCs were detached with Accutase, plated at 20,000 cells per well in a 96-well low-adherence plate in yolk sac induction medium plus the rock inhibitor Y-27632 (10 µM), and centrifuged at 1000 rpm for 3 min to aggregate the cells for embryoid body formation. Yolk-sac induction medium consisted of MAC iPS-Brew XF complete media plus bone morphogenetic protein 4 (BMP4, 50 ng/ml), vascular endothelial growth factor (VEGF, 50 ng/ml), stem cell factor (SCF, 20 ng/ml), and ß-mercaptoethanol. Yolk-sac induction media was given for the first 5 days of culture. On day 6, YS-EBs were plated in 100 mm cell culture plates in factory media consisting of XVIVO15 supplemented with 0.5% NEAA, ß- mercaptoethanol, macrophage colony stimulating factor (M-CSF, 100 ng/ml) and interleukin-3 (25 ng/ml). At 2-4 weeks after plating, human PMPs egress from the yolk sac lumen into the surrounding media and are continuously produced for more than 3 months.

To mature PMPs into microglia, PMPs were transitioned to maturation media composed of XVIVO15 supplemented with N2, Glutamax, 0.5% NEAA, interleukin-34 (IL-34, 100 ng/ml), granulocyte macrophage colony stimulating factor (GM-CSF, 10 ng/ml, and M-CSF (25 ng/ml) for 1-2 weeks. The medium was changed every 4 days. Cells were then detached with 20 min Accutase treatment and used for experiments.

### Generation of iPSC-derived Cerebral Organoids

Cerebral organoids were generated from 1 iPSC line using adapted previously published protocols (Qian et al. 2020). In this study, 9,000 iPSCs were cultured in low-adherence 96-well plates to form uniform embryoid bodies. On day 1, the medium was changed to neural induction medium composed of DMEM/F12 supplemented with 1X N2, 2 µM A83-01, and 2 µM dorsomorphin. On days 5 and 6, half of the medium was replaced with DMEM/F12 supplemented with 1X N2, 10 µM SB431542, and 1 µM CHIR99021. On day 7, EBs were embedded in a Growth-Factor-Reduced Matrigel ’cookie’ as previously described (Qian et al. 2020). Half of the medium was changed every two days until day 14. On day 14, organoids were dissociated from the Matrigel cookie and placed in low-adherence 6-well plates on an orbital shaker at 85 rpm. On day 16, organoids were fed 1:1 DMEF12: Neurobasal, 1X N2, 1X B27-RA, 0.5% NEAA, and 200 µM L-AA. Media was replenished every 2-3 days until day 30. On day 30, the medium was changed to 1:1 DMEF12: Neurobasal, 1X N2, 1X B27, 0.5% NEAA, 200 µM L-AA, and.25% GFR-Matrigel. Half of the media was changed every 3-4 days.

### Organoid Slicing

To create sliced neocortical organoid cultures, cerebral organoids were sectioned with a vibratome and transferred back into in vitro cell culture. Briefly, a fresh stock of 3% low-melting temperature agarose in PBS was prepared and kept at 39°C to remain liquid. Artificial cerebral spinal fluid (ACSF) for sectioning contained (in mM): NaCl 125, KCl 2.5, NaH_2_PO_4_·H_2_O 1.25, NaHCO_3_ 25, MgCl_2_ 1.2, CaCl_2_ 2.5, glucose (C_6_H_12_O_6_) 2.5, and sucrose (C_12_H_22_O_11_) 22.5. The ACSF was then bubbled with 5% CO_2_/95% O_2_ for 10 minutes before bivalent cations (in mM): MgCl_2_ 1.2, CaCl_2_ 0.625. The solution was then filtered through a 0.2-micron PES filter and returned to bubbling on ice. Cerebral organoids were then transferred to 37°C DMEF12, 200 µM L-AA, and 55 µM beta-mercaptoethanol before being embedded in 3% low-melting temperature agarose. The agarose block containing organoids was then sectioned at 400 µM thickness. Slices were then subjected to two washing processes in Primocin-treated media to reduce the chance of contamination.

### HIV-1 Stock Preparation

HIV-1 JRFL virus (ARP-395) and HIV-1 YU2 proviral DNA (ARP-1350) were obtained through the NIH HIV Reagent Program, Division of AIDS, NIAID. HIV-1 JRFL was propagated in phytohemagglutinin (PHA, 5 µg/ml)-stimulated human peripheral blood mononuclear cells.

Viruses were harvested on days 7-10 post infection. HIV-1 YU2 stocks were generated by transfection of 293T cells with HIV-1 YU2 proviral DNA. The transfected cells were cocultivated with PHA-PBMCs 48 h after transfection, and cell supernatants were harvested 7 days after coculture. The YU2 HIV stocks were further propagated in PHA-PBMCs as described. The amount of viral p24 was determined using the HIV-1 Gag p24 DuoSet ELISA (R &D System, DY7360-05).

### HIV-1 infection

PMPs were matured for 1-2 weeks in microglia maturation medium. Matured PMPs were then incubated with 2 ng/ml p24 of either JRFL strain or YU2 strain HIV-1 (NIH AIDS Repository) for 24 hours at 300,000 cells per 100 µl of media. After 24 hours, DMEF12 was added, and the cells were centrifuged for 3 min at 120 rcf. The cells were then resuspended in microglia maturation media minus M-CSF and plated. Half of the media was changed every 3-6 days.

### HIV-1 2-LTR circles PCR

For detection of HIV 2LTR circles, total DNA (including small nongenomic DNA) was isolated with a DNeasy Blood and Tissue kit (Qiagen 69504). End point PCR analysis (Taq 2X Master Mix, NEB) was performed using 300 ng DNA template and 2-LTR circle primers: forward, MH535: 5′-AACTAGGGAACCCACTGCTTAAG-3′; reverse, MH536: 5′-TCCACAGATCAAGGATATCTTGTC-3′ (Butler, Hansen, and Bushman 2001).

### Tissue preparation

Organoids were fixed in 4% PFA overnight at 4°C. Organoids were then transferred to 15% sucrose and 0.05% sodium azide in PBS for 24 hours, followed by another 24 hours in 30% sucrose and 0.05% sodium azide in PBS. Organoids were then embedded in OCT freezing medium and snap frozen on an ethanol dry ice slurry. Frozen embedded organoids were given at least 3 hours to warm to -20°C before cryosectioning on a Leica cryostat at thicknesses between 10-30 µM. Sections were mounted on charged microscope slides (VWR) and placed on a slide warmer at 37°C for 30 min. Slides were then stored in a slide box at -20°C until used for staining.

### Immunofluorescence

For immunocytochemistry, half of the medium was left in each well with an equal amount of 4% PFA and then added at a final concentration of 2% PFA overnight at 4°C. The next day, PFA was removed, and the cells were washed with PBS plus 0.3 M glycine for 30 min. The cells were then permeabilized with 0.2% Triton X-100 in PBS for 7 min. The cells were then blocked for 1 hour in blocking buffer consisting of 5% normal goat serum (NGS), 4% bovine serum albumin (BSA), 0.05% Triton X-100, 0.05% sodium azide, and PBS.

For immunohistochemistry, cryosectioned organoid slides were warmed to 37°C for 5 min. A hydrophobic barrier was then drawn around the specimens and allowed to dry for 5 min. PBS plus 0.3 M glycine was then added in a liquid dome over the specimens to rehydrate for 10 min. Sections were permeabilized with 0.5% Triton X-100, 0.3 M glycine, and PBS for 1 hr. After permeabilization, the sections were blocked with blocking solution consisting of 5% normal goat serum, 4% BSA, 0.05% Triton X-100, 0.05% sodium azide, and PBS for 1 hr. Primary antibodies were diluted in blocking solution and applied to the sections overnight at 4°C. The primary antibodies and their dilutions are summarized in the Key Resources Table. After washing with 0.05% Triton X-100 and PBS (PBST) 5 times, secondary antibodies diluted in blocking buffer were added for 1 hr at RT. Finally, the sections were washed five times with PBST before a final wash in deionized water and mounting with DAPI Fluoroshield (Fisher). Secondary antibodies were Alexa Fluor 488-, 546-, 633- or 647-conjugated goat antibodies (Invitrogen) used at a 1:500 dilution. Images were taken on a Zeiss LSM700 confocal microscope.

### RNA isolation

Total RNA was prepared with an RNeasy Plus kit (Qiagen) with preprocessing through Qiashredder (Qiagen) and frozen at -80°C.

### RT-qPCR

cDNA was generated using SuperScript VILO Master mix. RNA expression was measured using SYBR green on a QuantStudio 3 (Agilent). Probes were generated using NCBI primer blast and are listed in Table X. Two technical replicates were performed per sample. When a high standard deviation between replicates occurred, replicates were rerun in triplicate. All expression levels were normalized to GAPDH expression and represented as 2^^-(ΔΔCT)^. For null result values where a ct was not achieved, a max ct value of 40 was used for statistical analysis.

### Bulk RNA sequencing and analysis

We performed bulk RNA-sequencing analysis of iPSC-derived microglia. Total RNA was prepared using an RNeasy kit (QIAGEN). Library construction and sequencing were performed by Novogene. The libraries were subjected to 75 bp paired read sequencing using a NextSeq500 Illumina sequencer. Approximately 30–36 million paired reads were generated for each sample. Bc12Fastq software, version 1.8.4, was used to generate Fastq files. The genome sequence was then indexed using the rsem-prepare-reference command.

Each fastq file was trimmed and checked for quality with fastp (v. 0.12.2) and then aligned to the UCSC hg38 human genome using HISAT2 (v.2.1.0)71,72. Transcript counts were extracted using the featureCounts function of the Rsubread package.

### Statistics and reproducibility

All data represent the mean ± s.e.m. When only two independent groups were compared, significance was determined by a two-tailed unpaired t test with Welch’s correction. A p value <0.05 was considered significant. The analyses were performed in GraphPad Prism v.9. All experiments were independently performed at least three times with similar results.

---

**Figure S1:**
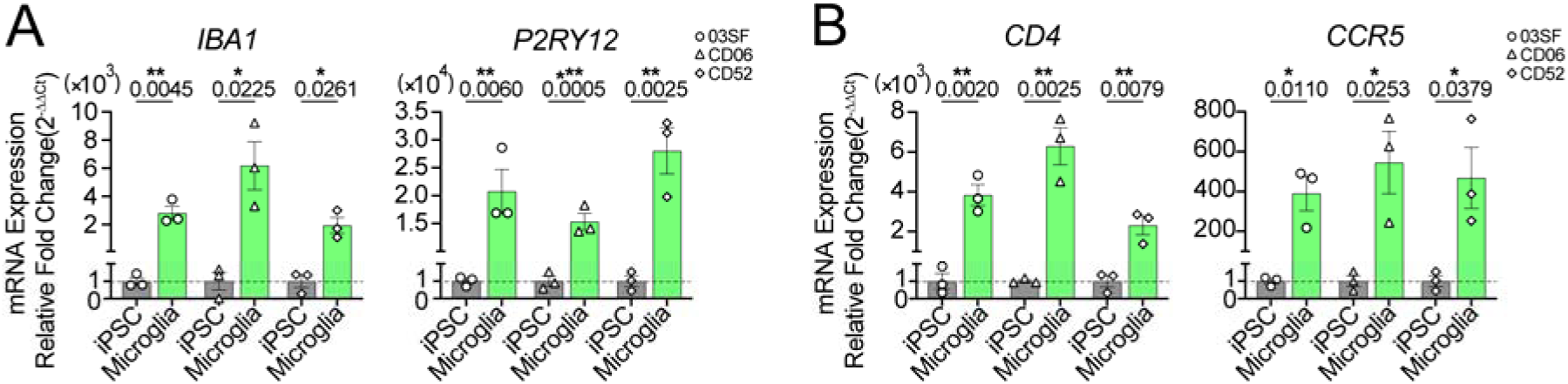
Microglial RT-qPCR analysis. (A) RT-qPCR analysis of *IBA1* and *P2RY12* mRNA expression in iPSCs and iPSC-derived microglia; n=3 cell lines (03SF, CD06, CD52) and 3 biological replicates/line; t test; data are presented as the mean ± SEM. (B) RT-qPCR analysis of *CD4* and *CCR5* mRNA expression in iPSCs and iPSC-derived microglia; n=3 cell lines (03SF, CD06, CD52) and 3 biological replicates/line; t test; data are presented as the mean ± SEM.

**Figure S2:**
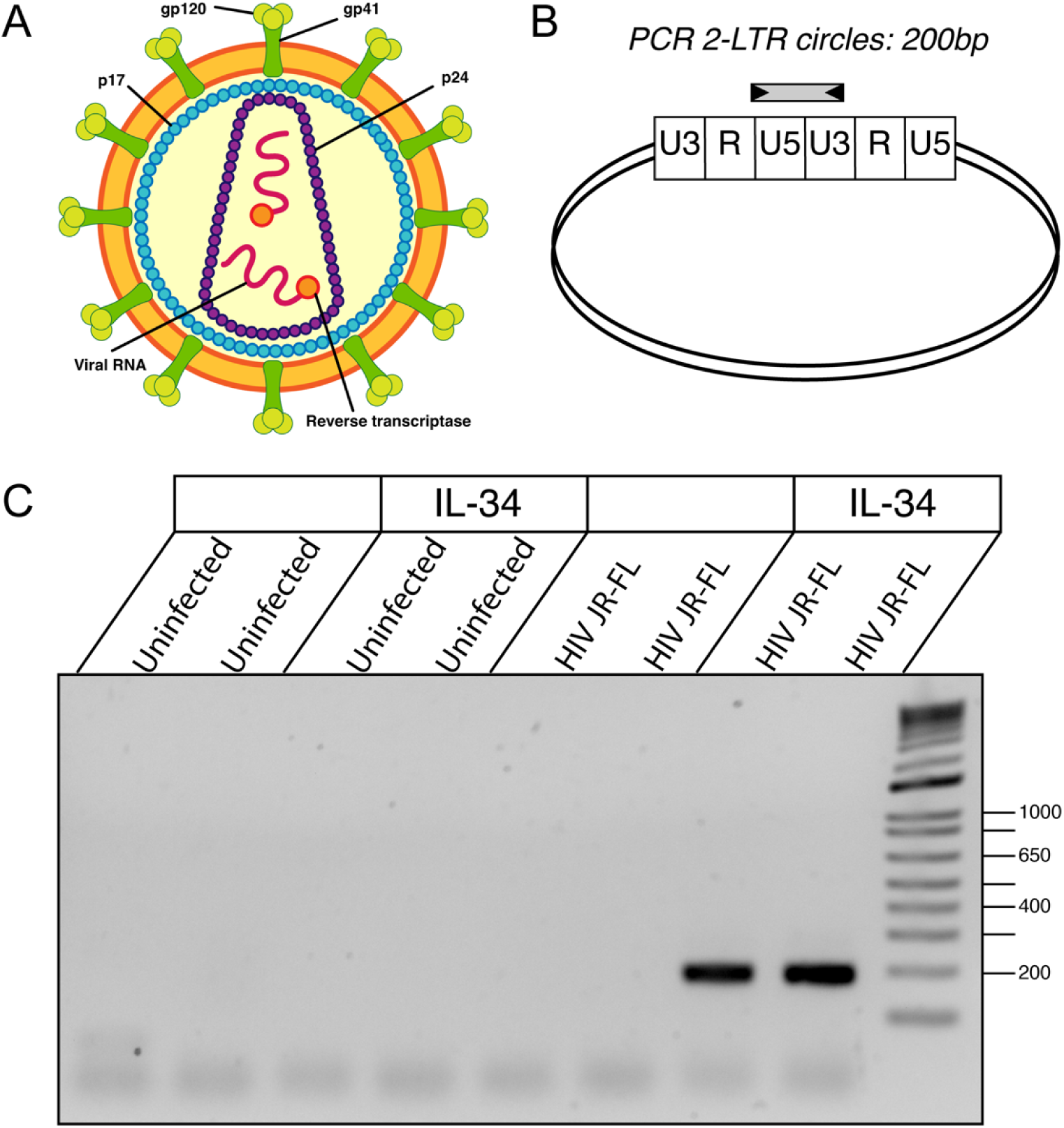
PCR analysis of 2-LTR circles. (A) A schematic representation of HIV-1 structure. (B) A schematic representation of an HIV-1 2-LTR circle. (C) PCR analysis of 2-LTR circles on day 4 post HIV-1 JRFL infection and mock infection.

**Figure S3:**
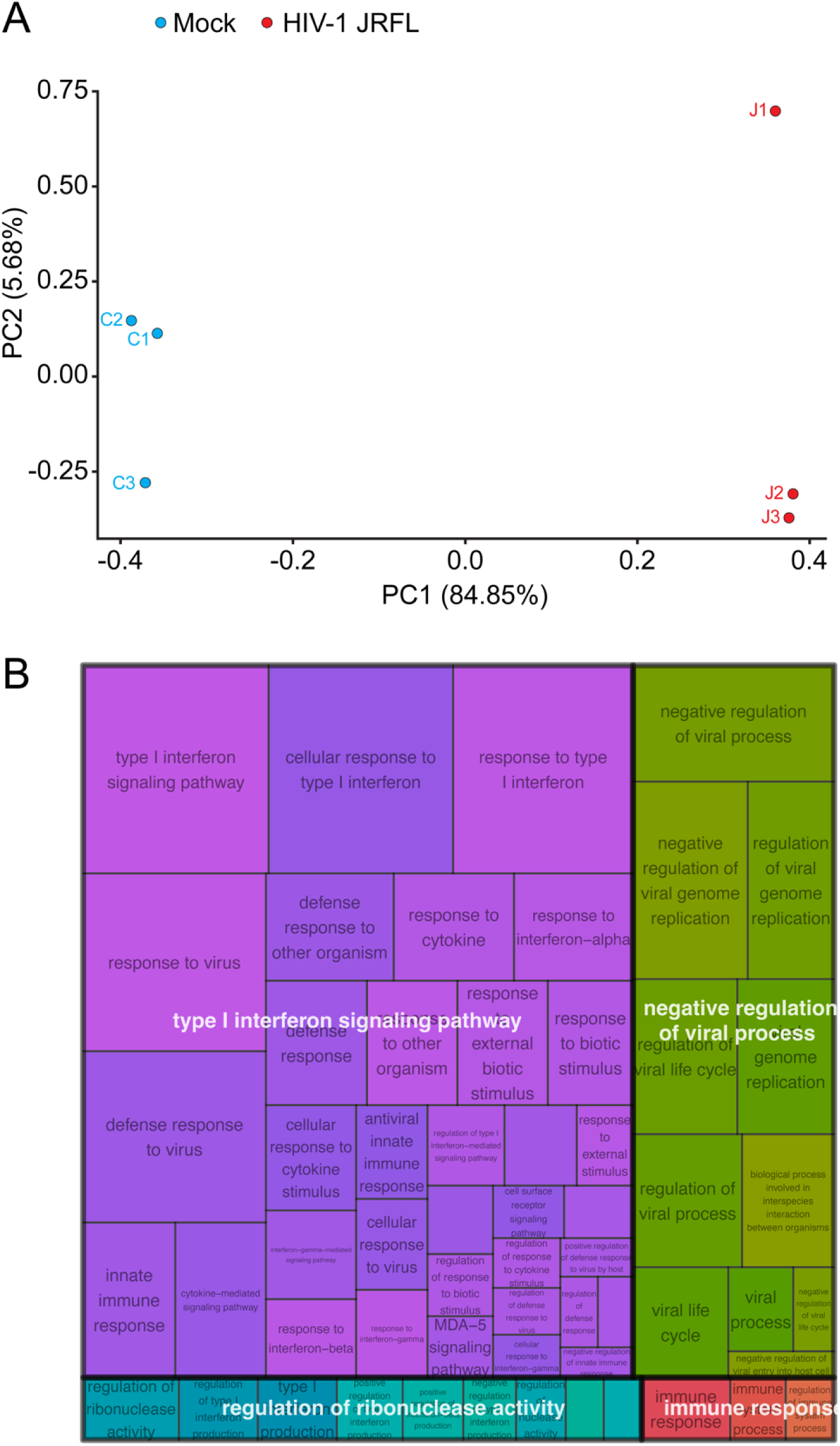
RNA-seq analysis of HIV-1 JRFL- vs mock-infected 03SF microglia. (A) A PCA plot of RNA-seq data collected from HIV-1 JRFL-infected microglia vs mock-infected microglia on day 16; n=3 independent infections of 03SF microglia. (B) A REVIGO treemap indicating major dysregulated pathways from gene ontology (GO) analysis of HIV-1 JRFL-infected microglia vs mock-infected microglia on day 16; n=3 independent infections of 03SF microglia.

